# Entrainment and maintenance of an internal metronome in premotor cortex

**DOI:** 10.1101/394783

**Authors:** J Cadena-Valencia, O García-Garibay, H Merchant, M Jazayeri, V de Lafuente

**Affiliations:** Institute of Neurobiology, National Autonomous University of Mexico. Boulevard Juriquilla 3001, Querétaro, QRO., México, 76230; McGovern Institute for Brain Research, Massachusetts Institute of Technology, Cambridge, MA, USA

**Author notes:** Corresponding author: Victor de Lafuente.

**Keywords:** Rhythm perception, supplementary motor area, local field potential, non-human primates, timing

## Abstract

To prepare timely motor actions we constantly predict future events. Regularly repeating events are often perceived as a rhythm to which we can readily synchronize our movements, just as in dancing to music. However, the neuronal mechanisms underlying the capacity to encode and maintain rhythms are not understood. We trained nonhuman primates to maintain the rhythm of a visual metronome of different tempos and then we recorded neural activity in the supplementary motor area (SMA). SMA exhibited rhythmic bursts of gamma band (30-40 Hz) reflecting an internal tempo that matched the extinguished visual metronome. Moreover, gamma amplitude increased throughout the trial and provided an estimate of total elapsed time. Notably, the timing and amplitude of gamma bursts reflected systematic timing biases and errors in the behavioral responses. Our results indicate that premotor areas use dynamic motor plans to encode a metronome for rhythms and a stopwatch for total elapsed time.

## Introduction

Adaptive behavior benefits from the ability to discern temporal regularities in the environment. To exploit these regularities, the brain must be able to measure time intervals between repetitive events (Buhusi and Meck 2005; de Lafuente, Jazayeri, and Shadlen 2015; Confais et al. 2012; Leon and Shadlen 2003), and use this timing information to anticipate future events (Goel and Buonomano 2014; Jazayeri and Shadlen 2010; Uematsu, Ohmae, and Tanaka 2017). This behavior is evident when we dance to music, which requires perceiving rhythms and generating movements in sync with them (Levitin, Grahn, and London 2018). Nonhuman primates and other vertebrates are capable synchronizing their movements to periodic rhythms (Merchant et al. 2013; Takeya et al. 2017), and we recently showed that monkeys can internally maintain rhythms of different tempos in the absence of overt motor actions (García-Garibay et al. 2016). Ample evidence indicates that cortical and subcortical motor circuits participate in behavioral tasks that require time perception and temporally precise behavioral responses (Mita et al. 2009; Crowe et al. 2014; Bartolo, Prado, and Merchant 2014; Merchant and Averbeck 2017; Grahn and Brett 2007; Ivry 2004; Murray et al. 2014). Nonetheless the neuronal mechanisms that allow motor structures to encode rhythms of different tempos, in the absence of motor commands, are not yet completely understood.

We developed a novel visual metronome task in which nonhuman primates had to observe, and then internally maintain, a temporal rhythm defined by a *left-right* alternating visual stimulus. Crucially, subjects performed had to track the rhythm in the absence of overt movements (García-Garibay et al. 2016). By uncoupling rhythm encoding and maintenance from motor actions, we aimed to identify the mechanism that allows the brain to internally maintain rhythms of different tempos. While monkeys performed the task, we recorded the local field potentials (LFPs) and spiking activity of single neurons in the supplementary motor area (SMA) that has been implicated in timing and rhythm perception (Buzsáki, Anastassiou, and Koch 2012; Pesaran et al. 2002). Our results show that bursts of lower gamma band activity (30-40 Hz) reflect the internally maintained tempos by a simple mechanism: the intervals defining the rhythm are encoded by the periodic onset of gamma bursts. Moreover, increasing amplitudes of gamma bursts reflected an estimate of total elapsed time (i.e. the total time since the rhythm began). Importantly, gamma bursts encoded both rhythm and the total elapsed time in the absence of sensory stimulation and overt motor activity.

## Results

### Monkeys can perceive rhythms and maintain them internally

We trained two rhesus monkeys (*M*. *mulatta*) to perform a visual metronome task (Figure 1A). While maintaining eye and hand fixation over the screen, monkeys saw a visual stimulus that appeared on one side, switched to the other, and the back to the initial location. This alternating stimulus defined three *entrainment* intervals of an isochronous rhythm. On each trial the interval duration was pseudo-randomly chosen to be 500, 750, or 1000 ms. In this manner, animals were presented with a visual metronome whose tempo was changed on a trial-by-trial basis (Figure 1A).

**Figure 1.**
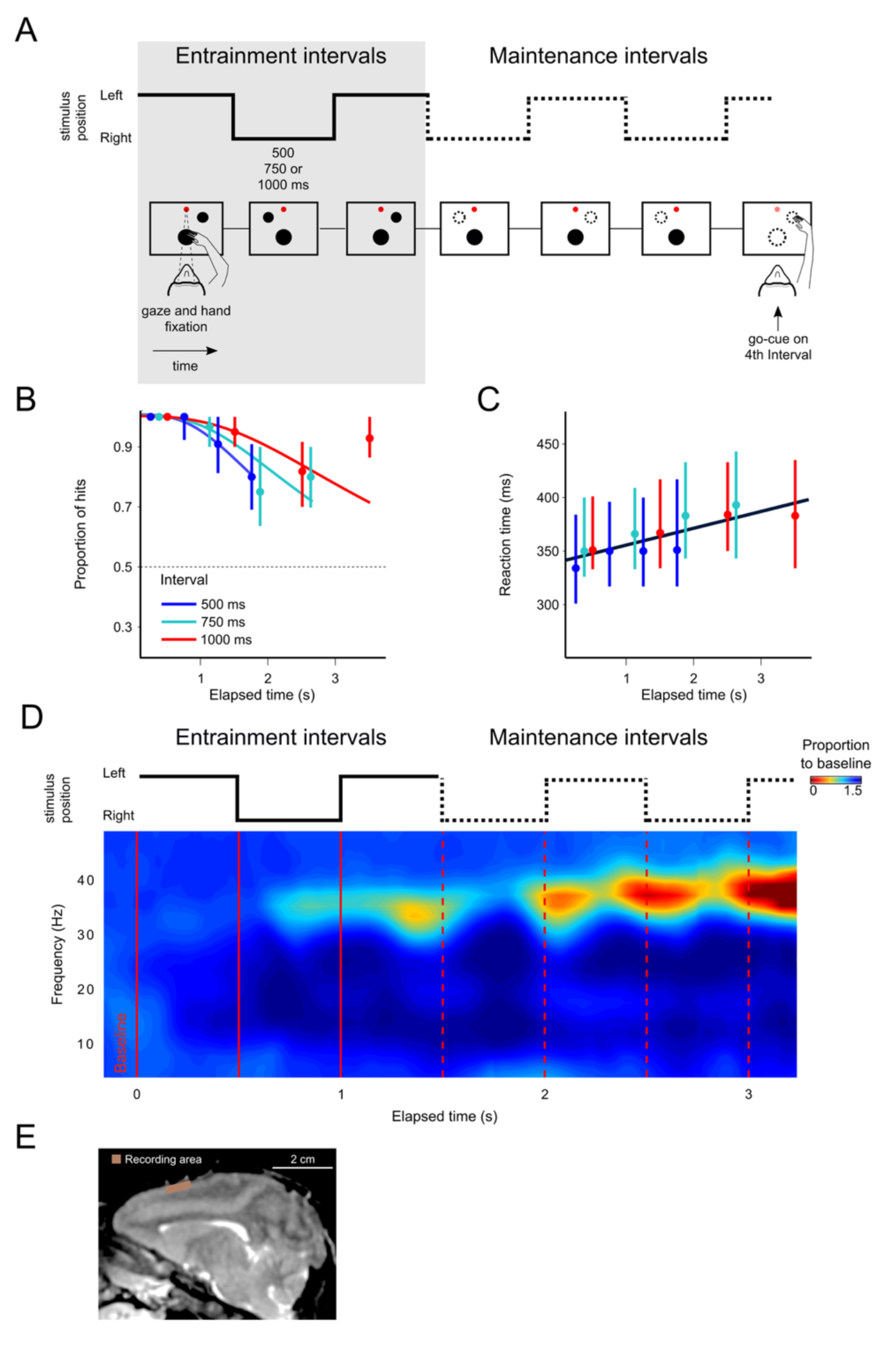
The visual metronome task. **(A)** Rhythms of different tempos were defined by a left-right alternating visual stimulus that appeared on a touch screen. While keeping eye and hand fixation, subjects first observed three isochronous *entrainment* intervals with duration of either 500, 750, or 1000 ms (pseudo-randomly selected on each trial). After the last *entrainment* interval, the visual stimulus disappeared initiating the *maintenance* intervals, during which the subjects had to keep track of the stimulus’ virtual location (left of right, broken lines). A *go-cue* (extinction of the hand fixation) at the middle of one of the four *maintenance* intervals prompted the subjects to reach towards the estimated location of the stimulus. It is important to note that this was not an interception task because the left-right switching stopped at the time of the *go-cue*. Monkeys received a liquid reward when correctly indicating the stimulus location. **(B)** The proportion of correct responses is plotted as a function of elapsed time during the *maintenance* intervals. Colors indicate the performance for the three tempos (500, 750, 1000 ms). Performance was significantly above chance (broken line at p=0.5; z-test p<0.001; n=131 sessions; median ± I.Q.R. over sessions). The decrease in performance as a function of elapsed time is expected from variability of the subjects’ internal timing in the absence of the external visual rhythm. This drop in performance was captured by a model of timing subject to scalar variability (continuous lines). **(C)** Reaction times to the *go-cue* increased as a function of elapsed time (n=131 sessions; median ± I.Q.R. over sessions). Black line indicates a linear regression on the median reaction times (dots). **(D)** Mean spectrogram across recording sessions and subjects (500 ms interval). The step traces at the top indicate the stimulus position as a function of time, for *entrainment* and *maintenance* intervals. Signal amplitude was normalized with respect to a 500 ms *baseline* period before stimulus presentation. A salient modulation of the LFP signal is observed around the gamma band (30-40 Hz). Gamma activity rhythmically increases in sync with the left-right transitions of the stimulus. Note also the increase in gamma activity as a function of total elapsed time. **(E)** Recordings were made from the medial premotor area, also called the supplementary motor area (SMA). The recoding chamber on monkey 1 (shown) was centered 23 mm anterior to Ear Bar Zero and 4 mm lateral to the midline, on the left hemisphere. The image shows a sagittal plane at 2 mm lateral from the middle.

After the third *entrainment* interval, the visual stimulus disappeared, and subjects had to maintain the rhythm internally by keeping track of the virtual position (left or right) of the stimulus as a function of elapsed time. To test the ability of subjects to maintain the rhythms, a *go-cue* at the middle of any one of up to four *maintenance* intervals instructed the subjects to reach towards the stimulus location (the *go-cue* consisted of removing the hand fixation point; the number of *maintenance* intervals was pseudo-randomly chosen; Figure 1A). Thus, the key parameters in the visual metronome task were (1) interval duration (500, 750, or 1000 ms), and (2) the number of *maintenance* intervals that subjects had to wait before the visual stimulus was gone.

We characterized monkeys’ ability to maintain the rhythms by plotting the proportion of correct responses as a function of the elapsed time since the initiation of the first *maintenance* interval (Figure 1B). The behavioral results show that monkeys satisfactorily performed the task and were able to correctly estimate the location of the stimulus in more than 80% of trials (94% ±0.2% monkey 1; 86% ±0.3% monkey 2; mean ± s.e. over sessions, n=131 sessions).

Importantly, performance as a function of time displays the hallmark of a timing task: the proportion of correct responses declines as a function of the number of *maintenance* intervals (or equivalently, elapsed time). The proportion of correct responses started close to 100% and declined to approximately 75% for the last *maintenance* intervals (last two data points for each curve). This behavior is consistent with the internal rhythm gradually drifting away from the true tempo of the stimulus (Gibbon et al. 1997; Grondin 2001). As we described in previous work (García-Garibay et al. 2016), this pattern is well captured by a model in which the subject’s time estimates arise from increasingly noisy (wider) distributions, described by Weber’s Law of time (also called the *scalar property* of timing) (Laje, Cheng, and Buonomano 2011). The increase in timing variability causes the subjects to eventually fall out of synchrony with the true stimulus position (getting ahead, or behind the true tempo), thus explaining the decrease in correct responses as a function of elapsed time (Figure 1B, the colored curves are fits of this model to the data; pooled data across monkeys).

Reaction times to the *go-cue* increases significantly in proportion to elapsed time within a narrow window ranging between 350 ms after the first maintenance interval of the fastest tempo (500 ms intervals) to 400 ms after the last interval of the slowest tempo (1000 ms intervals) (Figure 1C; R^2^=0.72, slope=11 ms/s, p<0.001; monkey 1 = 10.2 ms/s ± 0.8; monkey 2=11.3 ms/s ± 0.5). This increase in reaction times could be a result of the increasing difficulty in estimating the true stimulus position. As expected by scalar variability, the subject’s estimate of the stimulus position becomes noisier with time, thus increasing uncertainty and the reaction times necessary to make a decision. Overall, behavioral results show that monkeys were able to entrain to a rhythm, and maintain it in the absence of sensory stimuli, and importantly, in the absence of overt motor commands.

### Gamma oscillations reveal the internally maintained rhythms

While the monkeys performed the visual metronome task, we recorded neural activity in 131 experimental sessions (84 and 47 for monkeys 1 and 2, respectively; Figure 1E), and analyzed the local field potentials (LFPs) within 5-80 Hz band. As a first step we calculated the mean spectrogram for both monkeys, across all recording sessions (Figure 1D; 500 ms interval shown; combined data across monkeys). Modulations of LFP amplitude were especially salient in the 30-40 Hz frequencies, which we will refer to as gamma band. In this band, LFP power was up to two-fold larger than the baseline activity recorded 500 ms before trial initiation (p<0.001; permutation test of the time-frequency bins, 1000 permutations).

The LFP amplitude in the gamma band had a rhythmic structure. It increased markedly with the presentation of the last visible stimulus (3^rd^*entrainment* interval, Figure 1D), as well as near the time when the non-visible stimulus would be switching its position from one side of the screen to the other during *maintenance* intervals (Figure 1D; broken red lines). To test this observation quantitatively, we verified that the average gamma amplitude at the time of switches was significantly higher than halfway between them (t-test, p<0.01 for the three tempos; window sizes 1/4^th^ of interval length). In addition to the rhythmic modulation, gamma oscillations increased in amplitude as a function of total elapsed time (Figures 1D and 3C; note that the last *maintenance* interval displays the largest amplitude).

The analyses so far focused on mean LFP activity across sessions. To gain further insight into the LFP dynamics supporting the maintenance of internal rhythms, we analyzed LFP amplitude modulations within single trials. The LFP recordings from single trials (band-passed at 30-40 Hz) revealed short-duration bursts during which the oscillations transiently increase in amplitude (Figure 2B), consistent with recent findings in the prefrontal cortex (Lundqvist et al. 2016). Importantly, we observed that during the *maintenance* epoch, these bursts tended to coincide with the times at which the stimulus would have changed position, as is shown by the peaks in the spectrogram of the example single trials (Figure 2A).

**Figure 2.**
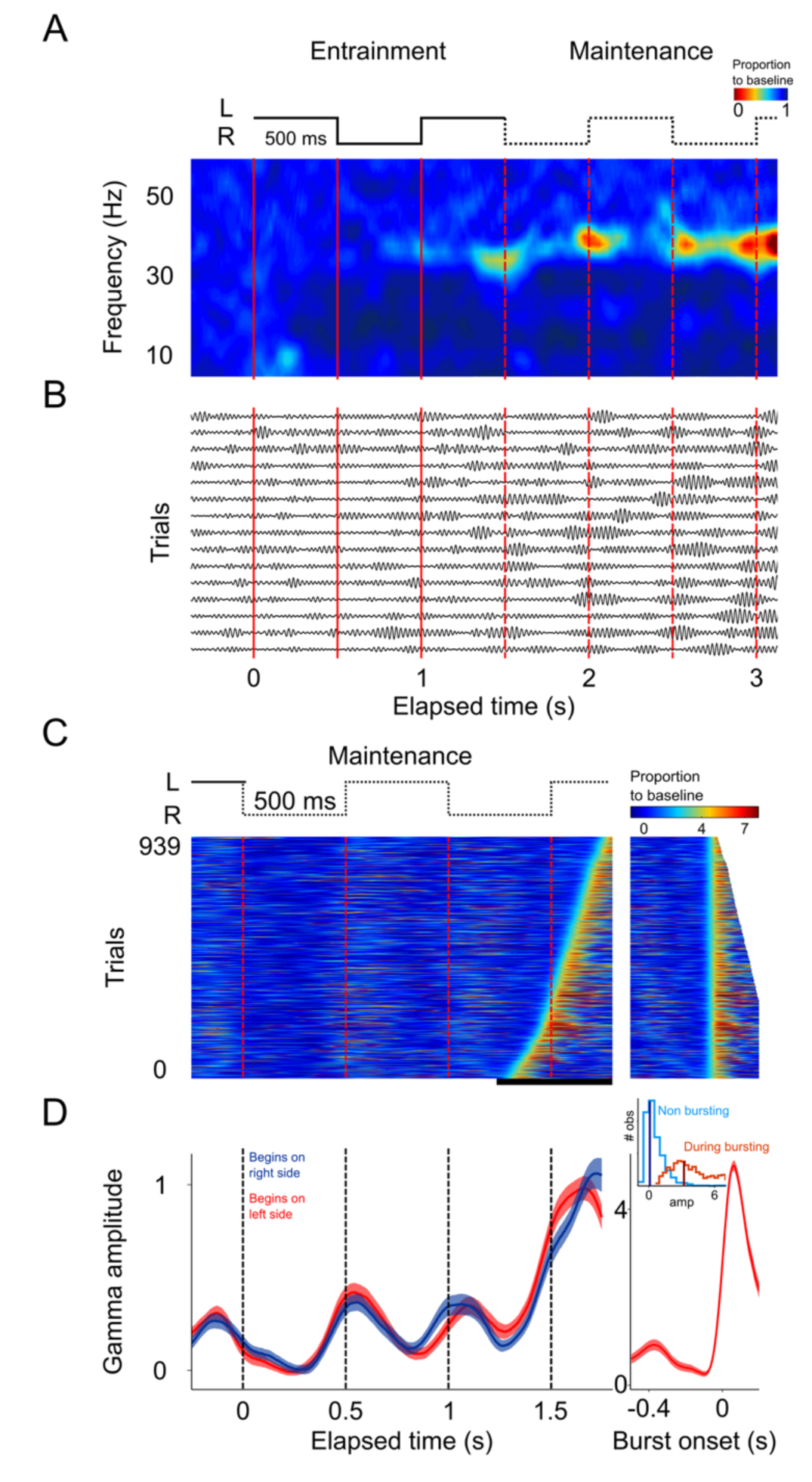
Single trial analysis of the LFP. **(A)** Spectrogram of 15 single trials of the 500 ms interval. There is an increase in amplitude at the gamma band (30-40 Hz) particularly salient during *maintenance* intervals. **(B)** Single-trials of the LFP signal, band-pass filtered at the 30-40 Hz gamma band. Gamma oscillations are composed of transient bursts during which oscillations increase in amplitude. Note how the bursts tend to occur in sync with left-right transitions of the stimulus and tend increase in amplitude as a function of total elapsed time. **(C)** Gamma amplitude on each trial is coded by color (939 trials; 500 ms interval). Trials were sorted according to burst onset time within the window marked by the black line at the bottom. The panel on the right shows the gamma bursts aligned to their onset time. Bursts were defined as the period in which gamma amplitude exceeded the 90^t^^h^ percentile of the amplitude distribution across trials, for at least 100 ms (4 cycles of the gamma rhythm). **(D)** Mean gamma amplitude is plotted as a function of elapsed time. Note how the periodic increases in gamma are in sync with the left-right internal rhythm during the *maintenance* intervals. The panel on the right shows the mean profile of the bursts in the last *maintenance* interval. The inset shows the distribution of the gamma amplitude during bursting (red distribution) and non-bursting (blue distribution) periods of the trials (dark vertical lines indicate the median burst amplitude for each distribution).

This trend is readily visualized by color-coding the amplitude of gamma oscillations and plotting all recorded trials on a single panel (Figure 2C). It is readily apparent that gamma bursts during *maintenance* tend to appear around the times at which the stimulus should be switching from one side of the screen to the other. This pattern is captured by the mean gamma amplitude, across trials, as a function of elapsed time (Figure 2D; p<0.01, t-test that compared amplitudes at the times of switch [0.5 and 1 s] versus amplitudes at the middle of the interval [0.75 and 1.25 s]; 125 ms windows).

These salient temporal features of the gamma LFP were consistent across the three interval durations (Figure 3A; 500, 750, and 1000 ms intervals). To better demonstrate the distribution of the gamma bursts, we sorted the trials according to burst onset time in each of the four *maintenance* intervals.

**Figure 3.**
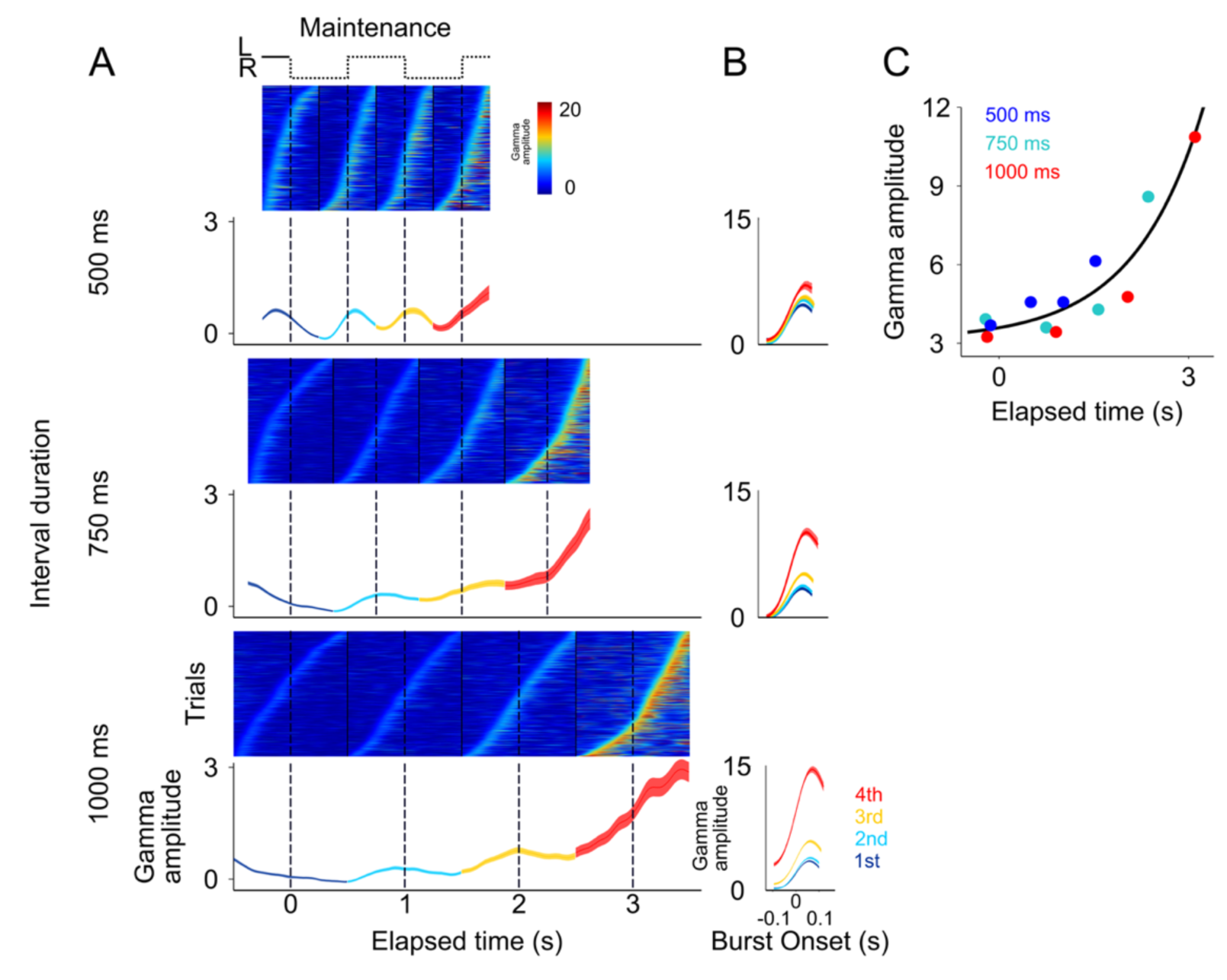
Gamma bursts in *maintenance* intervals for the three tempos (500, 750, 1000 ms). **(A)** For each interval and tempo, the bursts are ordered according to their onset time. Below single trials, the mean gamma amplitude is plotted as a function of time. **(B)** The temporal profile of gamma bursts is plotted for each stimulus transition (1^st^, 2^nd^, 3^rd^, and 4^th^), and for each interval duration (500, 750, and 1000 ms; top to bottom). The bursts have a stereotyped temporal shape and increase in amplitude after each consecutive transition. **(C)** Mean amplitude of gamma bursts plotted as a function of elapsed time for the three interval durations (500, 750, and 1000 ms).

It is important to emphasize that there are no motor actions during the *maintenance* intervals, and no periodic stimuli is shown on the screen. The only difference between the three groups of trials (500, 750, 1000 ms) is the tempo of the internal rhythm that subjects are maintaining. In other words, the rapid succession of the gamma bursts in the 500 ms intervals, and the more temporally distant bursts in the 1000 ms intervals, are a reflection of the subject’s internal maintenance of a visuo-spatial rhythm for the fast and slow tempos, respectively. This finding reveals a neural signature of rhythms, of different tempos, that are maintained internally.

Alignment of the gamma bursts to their onset time revealed that bursts have a similar temporal profile across tempos and elapsed intervals (Figure 3B). Importantly, we found that the amplitude of these bursts increased in proportion to the time elapsed since the initiation of the internal rhythm (Figure 3C, R^2^=0.86). The results presented so far indicate that (1) the LFPs in SMA encode internal rhythms by means of gamma bursts that occur in sync with the beats (i.e., location switch) of a visual metronome presented earlier; and that (2) these bursts increase in amplitude, providing a correlated for total elapsed time.

### Errors due to deviations of the internal rhythm from the objective tempo

In a previous study we demonstrated that human subjects tend to lag behind fast tempos and get ahead of slow ones (García-Garibay et al. 2016). This predicts that animals might systematically overestimate the 500 ms rhythms, and underestimate the 1000 ms rhythms. However, since animals only had two response options (left or right), it was not possible to use behavioral responses to disambiguate errors in which the animals were ahead or behind the true tempo. Nonetheless, we hypothesized that systematic over and underestimations of the intervals should be reflected in the patters of gamma activity in SMA. We therefore compared the profile of gamma activity as a function of time, on correct and incorrect trials (Figure 4A).

**Figure 4.**
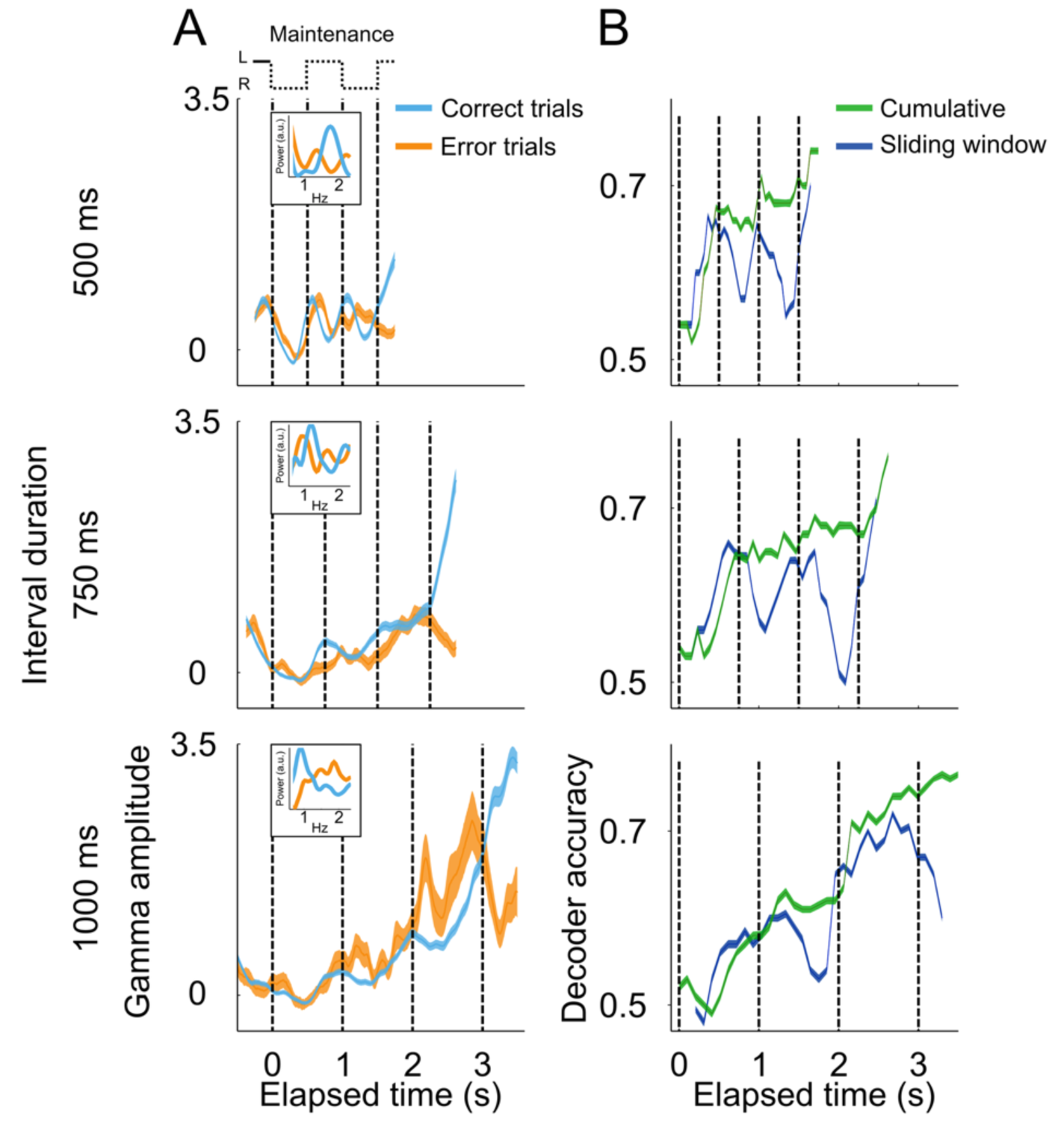
Gamma amplitude in correct and error trials. **(A)** Mean gamma amplitude during *maintenance* intervals, for correct (blue) and error trials (orange) (n=1400-3000 correct, 260-540 errors; colored area denotes s.e.m. across trials). The insets on each panel show the periodogram (power spectral density) of correct and error trials (mean across single trials). **(B)** Correct and error trials can be classified with increasing accuracy as a function of elapsed time. Two logistic classifiers were used to differentiate between correct and error trials (cross-validated on 50 correct and 50 error trials; n=100 iterations; line with shows s.e.m.). One used a growing window (cumulative, green line) that incorporated the gamma amplitude data as each trial developed, and the other used a sliding window of constant length across each trial (sliding window, blue line).

The results showed that, on the fast tempo trials (500 ms interval), the dynamics of gamma on error trials was right-shifted with respect to correct trials. That is, error trials displayed slower dynamics compared to correct trials (Figure 4A, upper panel). This trend was also captured by the power spectrums of error and correct trials, which showed that error trials indeed oscillated at lower frequencies (Figure 4A, inset on upper panel). Conversely, the dynamics of error trials on the slow tempo (1000 ms) resemble a left-shifted version of the correct trials, i.e., errors displayed faster dynamics as compared to the correct trials (Figure 4A; bottom panel). This pattern is also demonstrated by the power spectrums of correct and error trials, which show that error trials oscillated at higher frequencies compared to correct trials (Figure 4A, inset on the bottom panel). These results suggest that monkeys were lagging behind fast tempos and getting ahead of slow ones.

That the subject’s internal rhythm increasingly fell out of synchrony with the true tempo during error trials was also demonstrated by the ability of a logistic classifier (see Methods) to differentiate between correct and error trials, as a function of elapsed time (Figure 4B). This analysis shows that error trials are increasingly easier to decode as a function of elapsed time, just as it is expected from a rhythm that increasingly falls out of sync with the correct tempo. This pattern holds true for a classifier that cumulatively uses gamma amplitude information as the trial develops, and also for a classifier using the information from a sliding window as of constant length across the trial (Figure 4B).

### Gamma band activity in a *delayed-reach* task

Since SMA participates in the preparation of impending motor actions, it is possible that the rhythmic gamma bursts that we observed arise because this premotor area rhythmically prepare reach movements alternatively to the left and right locations of the screen. To test this possibility, we recorded the LFPs in a *delayed-reach* control task (Hwang and Andersen 2011) in which subjects were required to reach to the left or the right after being cued by a briefly presented visual stimulus (Figure 5A). In this task, monkeys waited a pseudo-randomly chosen time (1100 to 3000 ms, exponential distribution) before reaching towards the location specified by the cue (Figure 5B).

**Figure 5.**
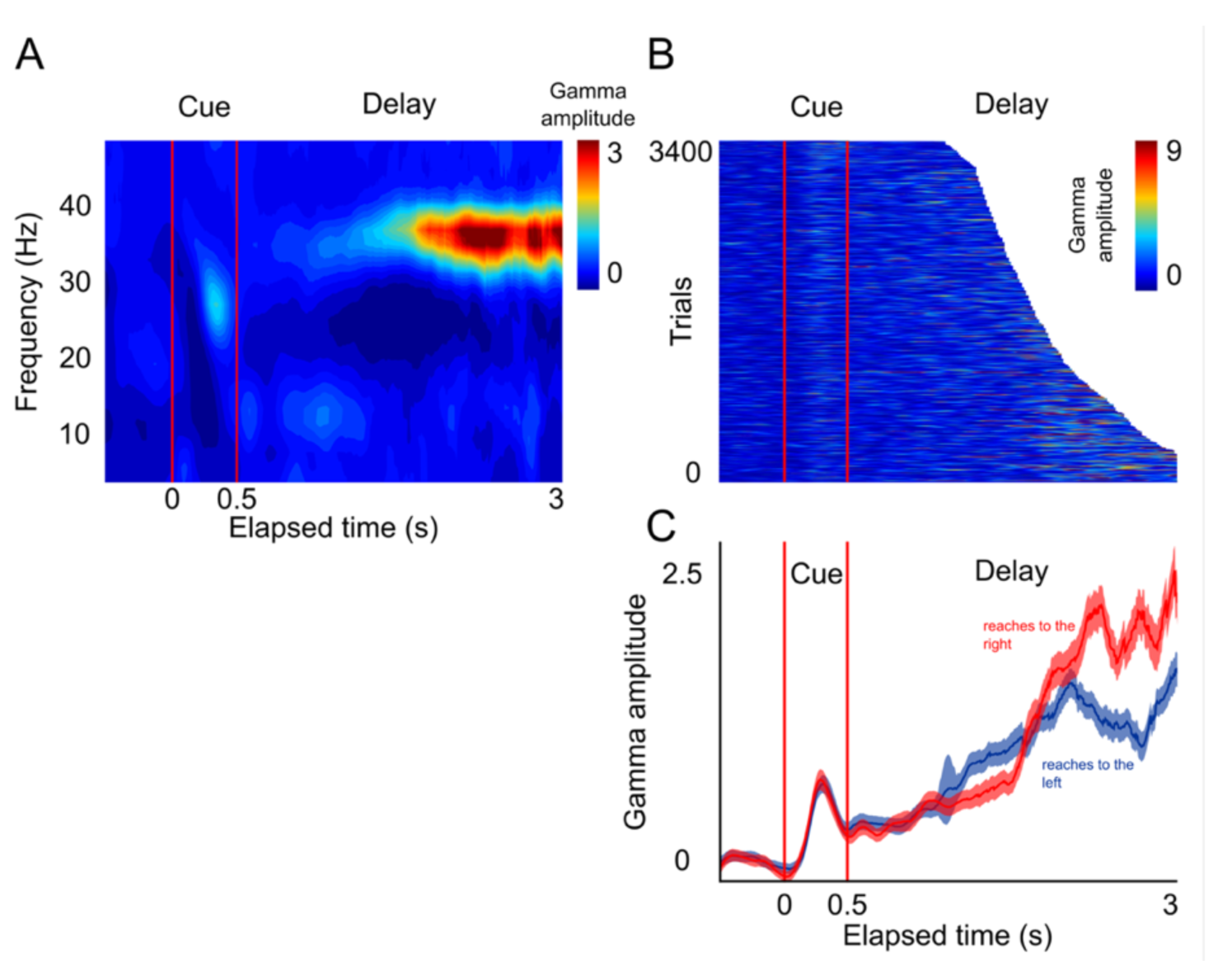
LFP activity in a *delayed-reach* task. **(A)** Mean spectrogram of the LFPs during the *delayed-reach* task (reaches to the right side of the screen are shown). The stimulus presentation is indicated by the red lines a 0-0.5 s (*cue*). After the sensory cue, a variable delay between followed (1.2-3 s; exponential distribution). A salient activation of the gamma band during the delay period can be observed. **(B)** Gamma amplitude across single trials of the delayed-reach task. **(C)** Mean gamma amplitude plotted as a function of elapsed time. After a brief sensory response, gamma activity increases as a function of elapsed time. Red and blue lines indicate reaches to the right and to the left, respectively.

The results of this control task show that, as monkeys prepare an impending reach movement, the LFPs in SMA generate bursts of gamma band activity that occur more frequently, and with increasing amplitude, as a function of total elapsed time (Figure 5C, *delay* period). These findings are consistent with the idea that gamma bursts in SMA encode impending motor commands. Moreover, the results of this control task are consistent with the idea that the SMA circuits reflect internal rhythms by means of rhythmically alternating motor plans to make a reach movement to the left and right locations of the screen.

### Gamma oscillations during entrainment of the visual metronome

According to the previous *delayed-reach* experiment, gamma bursts might be reflecting an internal rhythm by periodically alternating “*reach-left”* and “*reach-right”* motor plans. However, our task is designed such that a motor response was never required during the three *entrainment* intervals. For this reason, we next analyzed the gamma band activity during the *entrainment* intervals in which the presentation of the alternating visuo-spatial stimuli defined the different tempos of the visual metronome task (500, 750, and 1000 ms intervals; Figure 6A-B).

**Figure 6.**
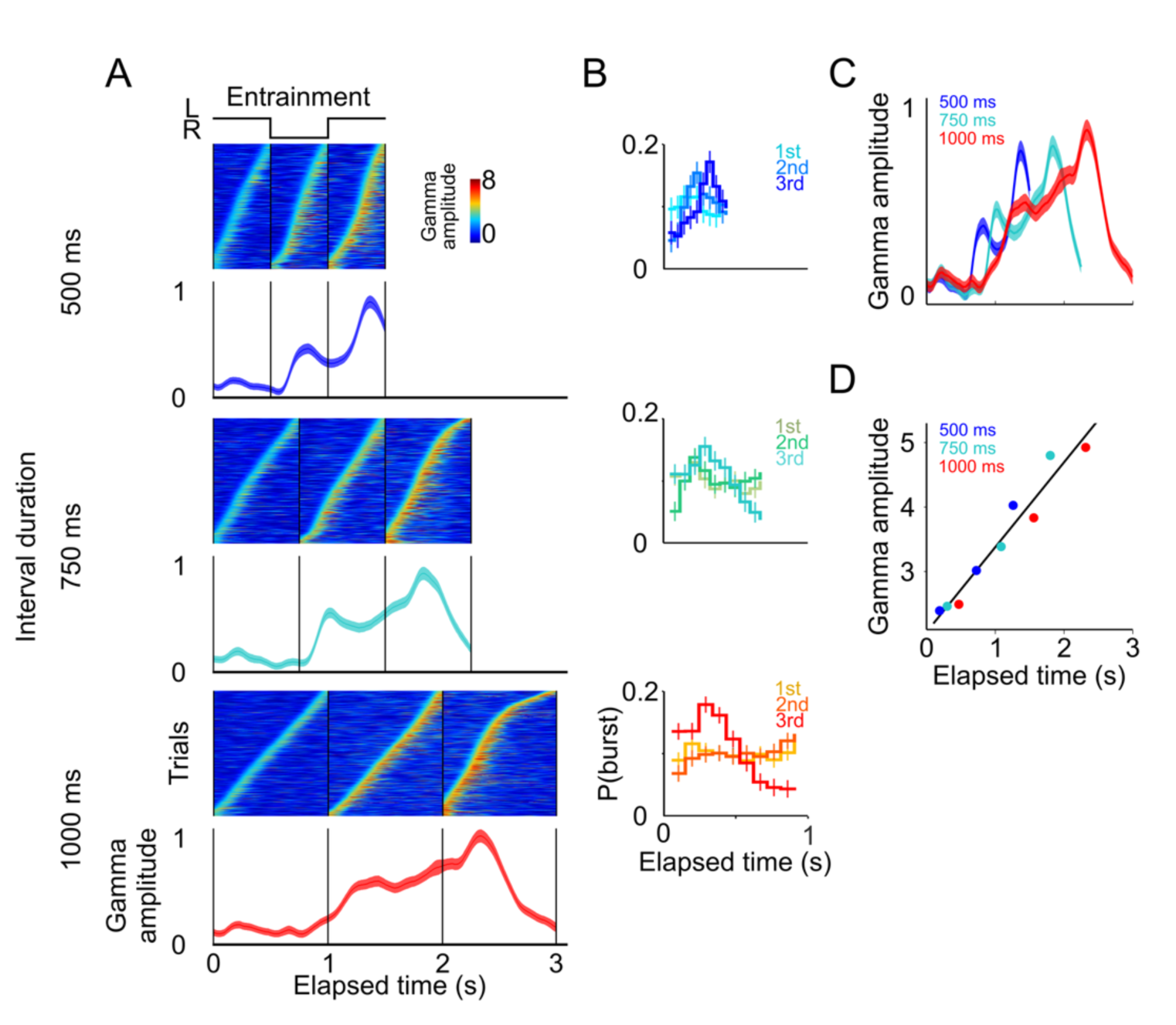
Gamma band activity in *entrainment* intervals. **(A)** Gamma bursts in single trials sorted by their onset time for each *entrainment* interval, and for each tempo (500, 750, and 1000 ms). Below the single trial panels, the mean gamma amplitude is plotted as a function of time. **(B)** Probability of burst onset plotted a function of elapsed time. Line color on each panel indicates the distribution of onset times for the consecutive *entrainment* intervals (1^st^, 2^nd^, and 3^rd^). **(C)** Mean gamma dynamics for the three tempos, plotted on the same timescale (same curves as the ones below single trial panels in (A). Note how mean gamma amplitude increases with each *entrainment* interval. **(D)** Burst amplitude as a function of elapsed time in *entrainment* intervals, for the three tempos.

The results showed that even during *entrainment* intervals, which did not involve any motor planning, bursts of gamma oscillations were present in each interval, and their amplitude progressively increased after the presentation of each visual stimulus (Figure 6A-C). It is important to note that gamma activity in *entrainment* intervals peaked after each stimulus presentation. This is in contrast to what was observed during *maintenance* intervals, in which the peaks of gamma occurred when the stimulus switched sides. We speculate that this phase offset could be related to the process of estimating interval duration, a process that necessarily happens during *entrainment* intervals.

A potential concern is that the gamma bursts in *entrainment* intervals are merely sensory responses to visual stimuli. However, a pure sensory response should produce similar gamma dynamics after each stimulus presentation, both across consecutive entrainment intervals (1^st^, 2^nd^, 3^rd^), and also similar across tempos (500, 750, 1000 ms), which was clearly not the case in our results (Figure 6A). In particular, two observations suggest that gamma bursts during entrainment cannot be explained solely in terms of a sensory response. First, gamma bursts increased in amplitude as a function of elapsed time, but the amplitude dropped sharply 500 ms after the onset of the third *entrainment* interval (Figure 6A, 750 and 1000 ms panels). Therefore, gamma bursts carry information about the animals’ knowledge that the third *entrainment* interval was the last visible interval, i.e. the last interval that could be used for estimating the tempo. Thus, gamma dynamics likely incorporate aspects of higher cognitive processing. Second, the times of burst onset do not have a fixed temporal profile with respect to stimulus presentation (Figure 6B). To demonstrate this, we measured the distribution of burst onset time across each consecutive interval (1^st^, 2^nd^, and 3^rd^ *entrainment* intervals) and across metronome tempos (500, 750, and 1000 ms), and then performed Chi-squared tests between these distributions (by using burst onset time we removed the effect of burst amplitude). The tests demonstrated that the temporal profiles of gamma onset times significantly differ, both across consecutive intervals and across metronome tempos (p<0.01; corrected for multiple comparisons). In fact, gamma responses to stimulus onset are similar only during first 500 ms of the first *entrainment* interval, which is the only epoch in which monkeys have no information about the metronome tempo (Figure 6C). These results indicate that gamma bursts reflect cognitive processes related to estimating the rhythm of the visual metronome.

To quantify the extent to which gamma burst amplitude encodes total elapsed time, we measured burst amplitudes in each of the three *entrainment* intervals (Figure 6D). We found that burst amplitude increased linearly in proportion to total elapsed time (R^2^=0.94). In this manner, in addition to periodically generating bursts in each *entrainment* interval, the SMA circuit reflected the total elapsed time since the beginning of the *entrainment* epoch.

### Neuronal spikes are associated with gamma band activity

Simultaneously with LFPs, we recorded the extracellular spike potentials of 113 neurons in the SMA (78 monkey 1; 35 monkey 2). The analysis of single unit activity will be the subject of a future report, but here we show five example units to illustrate the diversity of firing rate patterns in SMA during the visual metronome task (Figure 7).

**Figure 7.**
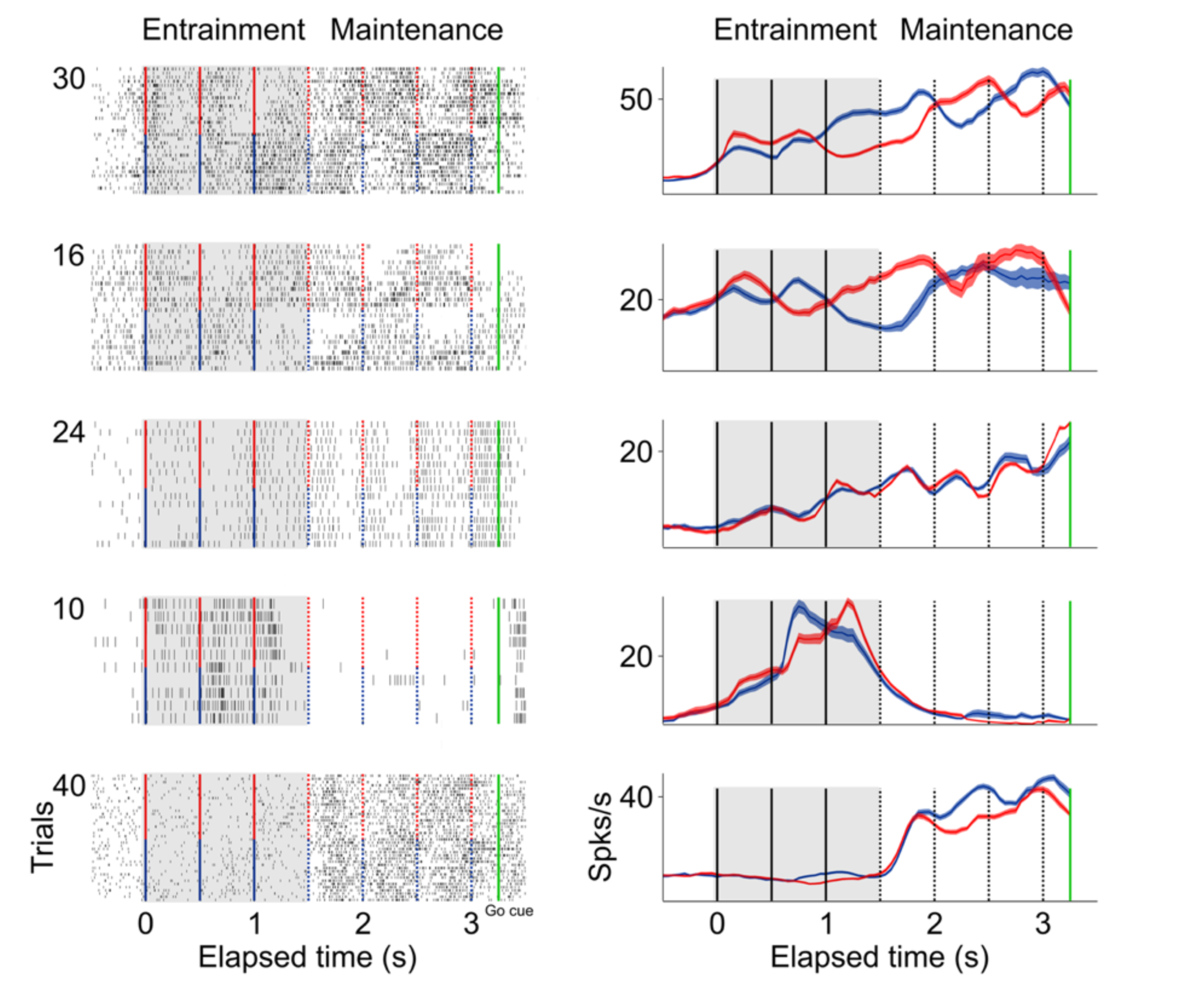
Firing patterns of single neurons during the visual metronome task. The panels on the left column show the raster plots of five example neurons during *entrainment* and *maintenance* intervals of the task. Green lines indicate the *go-cue* at the last *maintenance* interval. Red markers indicate intervals in trials were the stimulus initiated on the right, and blue markers when the stimulus initiated on the left. Panels on the right show the mean firing rate of each neuron. Firing patterns in SMA are diverse. The first two neurons (top to bottom) have preference for one side of the screen. The third neuron oscillates in sync with the stimulus tempo but shows no side preference. The fourth neuron is mostly active during the last two *entrainment* intervals. The fifth neuron is mostly active during the *maintenance* intervals.

To explore the relationship between single-neuron spiking and the simultaneously recorded LFP, we calculated the spike-triggered average (STA) LFP, and its spectral density, within a window of -100 to 100 ms surrounding each spike (Figure 8A-B, see Methods) (Denker et al. 2011; P. Fries 2001). We found that the LFP activity simultaneously recorded with each spike has a power peak at 30 Hz, and this peak is especially salient during *maintenance* intervals (Figure 8A, bottom panel; factorial ANOVA: interaction band/condition F=12.11 p<.05, Bonferroni tests of Gamma power in *maintenance* and *entrainment* vs baseline: p<.05). Moreover, the association between spikes and the 25-40 Hz frequency band is stronger at the times of stimulus transitions, i.e. around the times at which the stimulus switches from one side of the screen to the other (Figure 8C; window length around switch: half an interval, t test p<.005). To demonstrate that gamma is closely associated with the timing of spikes we performed a control analysis in which we jittered the spike times by ±15 ms with the resulting loss of the observed peak at the gamma band (random uniform distribution; grey traces Figure 8c; t test between jittered data in switch and non-switch conditions p=0.21).

**Figure 8.**
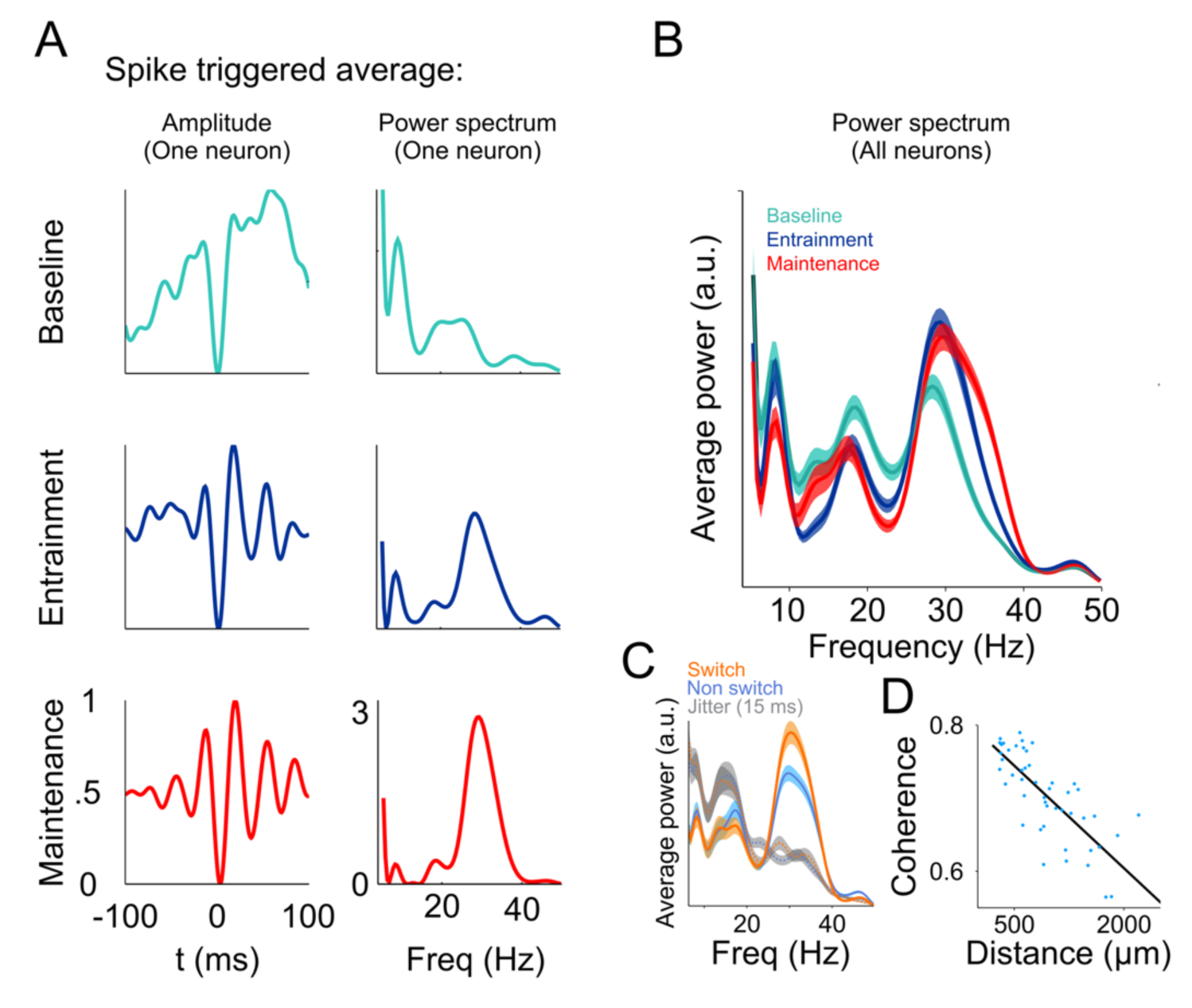
Relationship between spikes and LFP during the visual metronome task. **(A)** The three panels on the left show the spike-triggered average (STA) of the LFP signal surrounding each individual spike of an example neuron (-100 to 100 ms window centered at each spike time). The STA of three epochs is shown; *baseline* (cyan), *entrainment* (blue), and *maintenance* (red). The three panels on the left show the power spectrum of the STA on each epoch. **(B)** Average STA power across neurons. Colors denote trial epochs. Note the salient power of the STA around 30 Hz (colored areas show s.e.m. across neurons; n=113). **(C)** Average STA power for periods of stimulus switch and non-switch (windows of half the interval length, centered at times of switch or at the middle of each interval; colored areas show s.e.m. across neurons). Dotted lines and gray areas show the STA power obtained by jittering the spikes ±15 ms. **(D)** Coherence between the LFPs in simultaneously recorded electrodes, as a function of distance between them. The negative slope suggests that the recorded LFPs are generated by neuronal circuits in the vicinity of the recording electrodes.

These analyses demonstrate that the performance of the metronome task is accompanied by a tighter temporal relationship between the gamma bursts and the firing of single neurons, and this association is more prominent at the times of stimulus switching during the *maintenance* intervals. These results are consistent with previous investigations proposing that LFP oscillations near the gamma frequencies could help single neurons synchronize their firing, and thus have a larger and more temporally precise influence on downstream target structures (Siegle, Pritchett, and Moore 2014; Veit et al. 2017; Pascal Fries 2015).

Finally, we demonstrate that the LFP signals we recorded reflect local interaction and were not the result of signals being volume-conducted from other brain regions. We measured the coherence between LFPs of simultaneously recorded electrodes and plotted this measure as a function of the distance between them. The results show that coherence decayed as a function of electrode distance (Figure 8D, R^2^=0.72), as expected by an LFP signal that is generated in the neuronal circuits within the vicinity of the recording electrode.

## Discussion

Our results show that (1) monkeys can maintain rhythms in the absence of sensory stimuli and in the absence of overt motor commands. (2) Those internal rhythms are encoded by bursts of low gamma-band LFP oscillations in SMA whose timing and amplitude indicate rhythm intervals and total elapsed time, respectively. (3) The spikes of single neurons are associated with the low gamma band frequency of the LFP, which is consistent with the idea that gamma oscillations might help to synchronize populations of neurons whose temporally coincident firing would have a larger impact on its postsynaptic targets (Buzsáki 2015; Cardin et al. 2009; Pascal Fries 2015; Siegle, Pritchett, and Moore 2014; Veit et al. 2017; Womelsdorf et al. 2007; Wong et al. 2016).

The results of the *delayed-reach* task suggest that the bursts of gamma activity signal the preparation and impending execution of a motor plan (Merchant and Averbeck 2017; Mita et al. 2009; Chen, Scangos, and Stuphorn 2010; Ohara et al. 2001; Yokoyama, Nakayama, and Hoshi 2016; Shima and Tanji 2000). However, a purely motor explanation of our findings is not granted. First, gamma activity not only signals left-right transitions, but also total elapsed time. Second, bursts of gamma activity are also observed during the *entrainment* phase of the metronome task, a phase in which no motor actions are required. Instead, we favor the hypothesis that LFP signals in the premotor cortex are neither sensory nor motor, but encode a latent variable associated with a mental rhythm.

It has been debated whether subjects performing a rhythmic task time individual intervals, or instead rely on an estimate of total elapsed time (Laje, Cheng, and Buonomano 2011). Our results revealed that rhythms of different tempos are based on a neuronal representation of individual intervals, and there is also information available in the system about total elapsed time.

It is thought that gamma synchronization might be useful to the formation of local ensembles of neurons that increase the temporal coordination of presynaptic spikes on postsynaptic targets, allowing brief windows of effective communication (Wong et al. 2016; Womelsdorf et al. 2007; Buzsáki 2015). Previous results show that gamma oscillations increase before the execution of a motor action, and then shut down at the time of movement onset (Yokoyama, Nakayama, and Hoshi 2016) a result replicated by our data.

Previous work by Merchant and colleagues found that LFP gamma band activity in the basal ganglia was associated with the presentation of sensory stimuli defining the intervals within a hand tapping task (Bartolo, Prado, and Merchant 2014). They found that bursts of gamma were selective for intervals of different durations, and thus different cell populations were selective for different time intervals. We found no such duration selectivity in the SMA cortex, instead observing that gamma bursts encoded intervals of different durations.

Signals associated with timing tasks can be found across multiple brain areas, including parietal, motor, and premotor cortices, as well as dopaminergic midbrain neuron in the primate (Ghose and Maunsell 2002; Genovesio et al. 2006; Wise 2000; Mita et al. 2009; Harrington et al. 2010). For example, Jazayeri and Shadlen (2015) have shown that activity of single neurons in the lateral intraparietal area encodes the time elapsed time from a previous sensory stimuli, as well as the time remaining to initiate a saccadic eye movement (Jazayeri and Shadlen 2015). Importantly, they showed that these signals calibrate themselves according to the underlying probability to make an eye movement within a given temporal window. A recent important result by Jazayeri and colleagues demonstrated that encoding intervals of different lengths is achieved by means of speeding up or slowing down the temporal dynamics of populations of neurons that, individually, display widely different firing patterns (Wang et al. 2017). Our results extend this finding to the dynamics of the LFP oscillations by demonstrating that they also show temporal scaling (Figures 3A and 6C). A coherent picture is thus emerging, indicating that time-estimation and time-production signals are present as dynamic motor plans that are distributed across the motor structures that participate in executing timely motor actions.

## Acknowledgments

We thank Edgar Bolaños for technical assistance and Juan Ortiz for obtaining the MRI images. This work was supported by grants to from CONACYT Ciencia básica 254313 (VdL), 236836 (HM); Fronteras de la Ciencia 245 (VdL), 196 (HM); and PAPIIT IN207818 (VdL), IN202317 (HM). JCV is a doctoral student from Programa de Doctorado en Ciencias Biomédicas, Universidad Nacional Autónoma de México (UNAM) and received fellowship number 486768 from Consejo Nacional de Ciencia y Tecnología (CONACYT).

## Author contributions

JCV, OGG and VdL performed the experiments. VdL and MJ conceptualized the behavioral task. JCV performed the data analysis and generated the figures. All authors contributed to the interpretation of data. JCV and VdL wrote the first draft of the manuscript and all authors contributed to its final version.

## Declaration of interests

The authors declare no competing interests.

## Materials and Methods

### Subjects

Two adult male Rhesus monkeys (*Macaca mulatta*) participated in the study (weight: 5-7 kg, age: 5, 7 years). Experimental procedures were approved by the Ethics in Research Committee of the Institute of Neurobiology and were in agreement with the principles outlined in the Guide for Care and Use of Laboratory Animals (National Institutes of Health). Each monkey was surgically implanted with titanium head bolts and a titanium recording chamber over the left supplementary motor area (SMA). Placement of the chambers over the SMA was guided by structural MRI for both monkeys (Figure 1E).

### Behavioral Task

Monkeys were trained in a visual metronome task described in detail in a previous report (García-Garibay et al. 2016). Briefly, while maintain eye and hand fixation over a touch screen (ELO Touch Solutions, model 1939L; ASL Eye-Track 6), subjects observed a visual stimulus (gray circle, 10° diameter, 25° eccentricity) that periodically changed position from one side of the screen to the other, at regular intervals (*entrainment* epoch; 500, 750, or 1000 ms interval; pseudo-randomly selected on each trial; Figure 1A). After three *entrainment* intervals the visual stimulus disappeared, and subjects had to continue estimating its position (left or right) as a function of elapsed time (*maintenance* intervals; Figure 1A). This visuo-spatial rhythm task is similar to a visual metronome that paces a rhythm which subjects have to keep internally during the *maintenance* epoch. To quantify the ability of the subjects to maintain rhythms of different tempos a *go-cue* (disappearance of the hand fixation area) was presented at the middle of any of the four maintenance intervals (randomly selected, uniform distribution; Figure 1A). This *go-cue* instructed the subjects to make a reach movement towards the estimated target position (left or right). It is important to note that this was not an interception task, i.e., once the *go-cue* was presented the non-visible stimulus no longer changed position. Performance was measured as the proportion of correct responses plotted as a function of the elapsed time since the initiation of the *maintenance* epoch (Figure 1B). Visual stimuli and task control was achieved with the Expo software (designed by Peter Lennie, maintained by Robert Dotson; available at https://sites.google.com/a/nyu.edu/expo/).

### Neural Recordings

Neural recordings were performed with seven independent movable microelectrodes (2-3 MΩ, Thomas Recordings, Giessen, Germany). Electrodes were advanced in the coronal plane into the supplementary motor area until single unit activity was obtained in at least one of the electrodes. At each recording site, spikes were isolated online (Cerebus acquisition system, Blackrock Microsystems, Salt Lake City, UT, USA) and sampled at 30 KHz. The local field potentials (LFPs) were obtained by filtering the electrode signal at 0.5 to 500 Hz, at a 2 KHz rate. Offline, the signal was down sampled to 1 KHz, and band-pass filtered to the 2-50 Hz band.

### Data Analysis

Analyses were performed with MATLAB 2013b (The Mathworks, Natick, MA, USA) using custom code in combination with the Chronux Toolbox for the time-frequency maps (Partha P Mitra and Bokil 2007).

### Time-Frequency Decomposition

Spectral estimation was performed using multitaper methods (Pesaran et al. 2002; P.P. Mitra and Pesaran 1999; Cohen 2014). A 200 ms windows sliding at 5 ms steps was used for the time-frequency maps (one taper was used, 5 Hz bandwidth). Spectrogram power was normalized by dividing each frequency and time bin by the average power in a 500 ms *baseline* window before trial initiation.

### Single Trial Analysis

To characterize how the amplitude of the gamma oscillations is modulated over time, we averaged the normalized spectrograms over the low gamma band frequencies (30-40 Hz). Narrow-band filtering with analytic envelopes and complex Morlet wavelet convolution yielded similar results. Gamma bursts were defined as the period of time in which gamma amplitude exceeded the 90^th^ percentile of overall activity for at least 100 ms (i.e. for at least 4 cycles of the gamma oscillations).

### Classification of correct and error trials

A logistic function was used to identify correct and error trials:

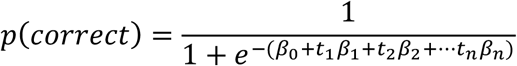

were t1 correspond to the gamma amplitude in the first time-bin, t2 to the amplitude on second time bin, and so on (10 time bins per interval, 35 time-bins for each trial). Thus, the predicted behavior arises from a linear combination of the gamma activity used to fit the logistic function. The classifier accuracy was measured on 100 trials (50 correct and 50 error trials; randomly selected) not used in fitting the logistic function. Fitting and testing was repeated 100 times, randomly selecting the test trials. For the cumulative window classifier (Figure 4B, green line) we used the gamma amplitude on the first time-bin and then tested the accuracy of decoding, then we added the data of the second time-bin and recalculated accuracy, and so on until the last time-bin. In a second approach that we called “sliding window”, a window of 5 time-bins were used to fit the classifier and calculate accuracy. This window moved across the trial to calculate accuracy as a function of elapsed time (Figure 4B, blue line).

### Spike-Triggered Average (STA)

To estimate the synchronization between the spikes and the simultaneously recorded LFP, 200 ms windows centered on each spike were analyzed (P. Fries 2001; Denker et al. 2011). The average LFP in these windows were computed and normalized peak-to-valley to values between 0 and 1. This procedure was applied before spectral decomposition of the STA (Figure 8A, power spectrum), allowing the comparison of spectral density maintaining the same maximum amplitude across conditions (*baseline, entrainment* and *maintenance* epochs). To assess statistical significance, we performed a factorial ANOVA with the factors *condition* (*baseline, entrainment, maintenance*), and *frequency* (alpha, beta, gamma), where the dependent variable was the average amplitude between 6 to 10 Hz for alpha, 15 to 24 Hz for beta and 30 to 40 Hz for gamma. This analysis demonstrated that the average power of the STA over the gamma band was significantly larger during *entrainment* and *maintenance*, as compared to the baseline period (p<0.01; Figure 8B). We normalized the amplitude of the LFP traces surrounding each spike to account for the increase in gamma amplitude with total elapsed time.

### Coherence between simultaneously recorded electrodes

To assess the locality of the observed LFP oscillations we estimated the phase clustering between the LFPs in pairs of simultaneously recorded electrodes. We used the time series of all trials recorded while the monkeys performed the task. For each electrode pair, we band-pass filtered the signal (30-40 Hz) and estimated the analytic envelope to obtain the instantaneous phase. Then, for each time point we estimated the difference angles between signals in the complex plane. The coherence was defined as the length of the average vector of all difference angles, a procedure that results in magnitudes between 1 (all difference angles are aligned to the same direction) and zero (random distribution) (Cohen 2014). To quantify how coherence decreased as a function of electrode separation we grouped the distance variable into 50 bins containing the same number of observations per bin. A linear regression was then applied to these data (Figure 8D).

